# Intra-male reduction in sperm size persists during growth in cuttlefish despite the absence of insemination-site dimorphism

**DOI:** 10.64898/2026.07.19.739393

**Authors:** Kyohei Kudamatsu, Noritaka Hirohashi

## Abstract

In certain groups of squids and cuttlefish, males display alternative reproductive tactics (ARTs). In squids, size-associated male dimorphism appears in mating posture, spermatophore transfer site (insemination dimorphism), and sperm flagellar length (sperm dimorphism). Sperm dimorphism is closely linked to insemination dimorphism, in which sperm are deposited either at the female’s external buccal mass or within her mantle cavity. Insemination dimorphism shapes post-copulatory sperm environments, including the mode of storage, the risk of sperm competition, and the site of fertilization. Therefore, sperm dimorphism is regarded as an adaptive consequence of insemination dimorphism. Conversely, in cuttlefish, both large consorts and small female-mimicking (sneaker) males deposit spermatophores in the same region of the female buccal mass, indicating the absence of insemination dimorphism. In *Sepia esculenta* and *Sepia lycidas*, sperm located at the distal end of the male reproductive tract possess longer flagella than those near the testis. This pattern is consistent across all male sizes, suggesting that as males grow, they produce sperm with shorter flagella. Additionally, these within-individual differences in sperm size are greater in smaller males and are positively correlated with relative testis mass. These findings indicate that insemination dimorphism is not required for the evolution of dimorphic sperm in cuttlefish. In both squids and cuttlefish, sperm from smaller males must either enter the female receptacle for extended storage or swim faster to compete with the more abundant consort sperm. These unfavorable conditions associated with the sneaker tactic may drive costly sperm evolution.

## Introduction

In certain groups of squids and cuttlefish, males display two or more discontinuous mating strategies, referred to as alternative reproductive tactics (ARTs), that are tightly linked to the male’s adult body size (Hanlon and Messenger, 2018). Large males (consorts) are predominant in acquiring mates and usually take an assertive approach to females, often after winning premating physical combat or visual display contests with rivals. Small males (sneakers) tend to be recessive in mate acquisition, leading to sneaky or early (long before egg spawning) copulations regardless of the presence or absence of a consort male (Apostolico and Marian, 2019; Naud and Havenhand, 2006; Naud et al., 2016). In other cases, the choice of male mating strategies is determined by their size relative to the female they are mating with (Wada et al., 2005a) or the timing of egg laying (Apostolico and Marian, 2019), and thus females also play an active role in selecting mating positions (Lin and Chiao, 2017). In any case, a male’s body size is the primary factor in determining a preferred mating strategy. In the family Loliginidae, male ARTs result in dimorphism in the site of spermatophore transfer to the female (insemination dimorphism), i.e., transferring at an internal location of the mantle cavity by consorts or at an external location of the buccal mass by sneakers (Fig. 1A) (Iwata et al., 2011). Previous studies with five species in the Loliginidae family have shown that the sperm from sneakers are generally longer than those of consorts (Apostólico and Marian, 2018a; Hirohashi et al., 2021; Iwata et al., 2011), suggesting that sneakers must produce more costly sperm than consorts to obtain fertilizations in competitive situations. Indeed, there are many differences in post-mating conditions between sneakers and consorts, such as the mode of sperm storage, intensity of sperm competition and the fertilization environment. Among these differences, we have long held the hypothesis that the presence of two separate insemination sites in females is a key factor contributing to sperm dimorphism. Insemination site dimorphism generates genuine differences in post-ejaculatory sperm fate, as described above. We therefore speculated that one or more of these distinct characteristics, resulting from insemination dimorphism, drive changes in sperm size to facilitate adaptation to different post-insemination environments.

**Fig. 1.**
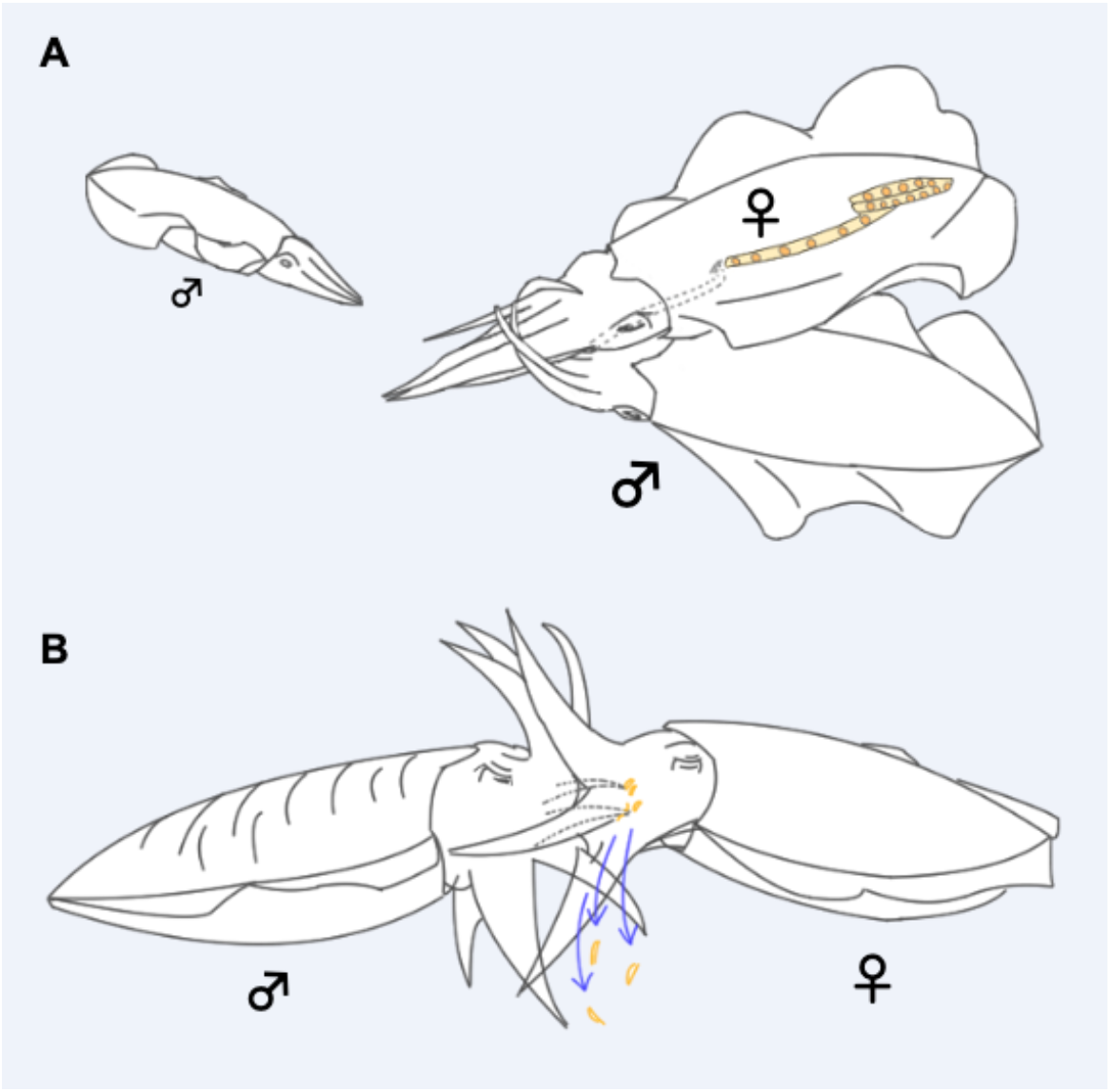
Mating strategies in some squids and cuttlefish. **A**, This illustrates alternative reproductive tactics in squid: a small sneaker male attempts copulation (*left*), while a large consort male (*right*) transfers his spermatophores to the oviductal end within the female’s mantle cavity. **B**, A male cuttlefish removes spermatangia deposited by other males from a female before transferring his own spermatophores. Schematic drawings were generated in accordance with the figures presented by Iwata et al. (2011) and Wada et al. (2005).

In cuttlefish, ARTs have also been reported in several species, including *Sepia plangon* (Brown et al., 2012), *Sepia apama* (Hanlon et al., 2005; Norman et al., 1999), *Sepia officinalis* (Cooke et al., 2017), and *Sepia pharaonis* (Lee et al., 2016). Smaller males employ a sneaky tactic to obtain copulations when a consort male is distracted by rivals, or disguise their body patterns to resemble females, thereby avoiding aggression from guarding males. This strategy is referred to as female mimicry (Hanlon et al., 2005; Norman et al., 1999). In *Sepia esculenta* and *Sepia lycidas*, there have been no field observations clearly documenting sneaky behaviors, however, in experimental settings in captivity, smaller males in both species exhibited copulations, suggesting that they are mating-competent in the wild (Wada et al., 2005b; Wada et al., 2010). Additionally, it has been recognized that males in these species, as well as in other Sepia species (Geary Boal, 1997; Hall and Hanlon, 2002; Naud et al., 2004), are capable of removing the existing spermatangia, transferred by previous males, from a mating female before attempting to transfer their own spermatophores (Fig. 1B). This behavior, collectively known as ‘sperm removal’, may occur as the consequences of polyandry and last male sperm precedence (Squires et al., 2015). Spermatangia are found attached exclusively to the limited area of the ventral side of the female buccal membrane, immediately adjacent to the seminal receptacle (SR) (Wada et al., 2005b; Wada et al., 2010). Under intense mating competition, males may prefer to transfer their spermatophores to this area because of its proximity to both the SR and the egg-deposition site, thereby achieving immediate fertilization success (Akter et al., 2026). Given the substantial variation in adult male size in cuttlefish, ranging from 185 to 310 mm in *S. lycidas* (Wada et al., 2010) and from 90 to over 200 mm in *Sepia officinalis* (Agus et al., 2024), as well as the likelihood of high polyandry inferred from complex behaviors such as female mimicry, sperm removal and narrowing of the insemination site, it is hypothesized that smaller males in *S. esculenta* and *S. lycidas* may engage in mating strategies that allow them to mate opportunistically when larger males are unaware. Taken together, it is of great interest to compare sperm size in cuttlefish between small sneakers and large consorts, because both types of males transfer sperm to the same area of the female; thus, no difference in the fertilization environment is expected (Wada et al., 2010). If insemination dimorphism is a prerequisite for the evolution of dimorphic sperm, then no difference in flagellar length would be expected to occur in cuttlefish or vice versa.

## Materials and methods

The cuttlefish, *S. esculenta* (Fig. 2A; 31 males, 25 females) and *S. lycidas* (Fig.2C; 15 males, 10 females) were obtained through shore jigging using standard squid fishing gear (Fig. 2A, *right*) from April 17 to June 11, 2025 (for a total of 51 days) at Senzaki Bay and Fukawa Bay in Yamaguchi, Japan. The squids were immediately anesthetized with seawater containing 3.75% magnesium chloride (Abbo et al., 2021), followed by cutting nerve cords between eyes. The cuttlefish specimens were measured (mostly on the same day of fishing) for dorsal mantle length, body weight, total gonad weight, testis weight, and spermatophoric complex weight. We calculated the gonadosomatic index (GSI*) and testicular somatic index (TSI*) using the following formulas.

GSI* = 100 x (total gonad weight) / ((body weight) - (total gonad weight))

TSI* = 100 x (testis weight) / ((body weight) - (total gonad weight))

**Fig. 2.**
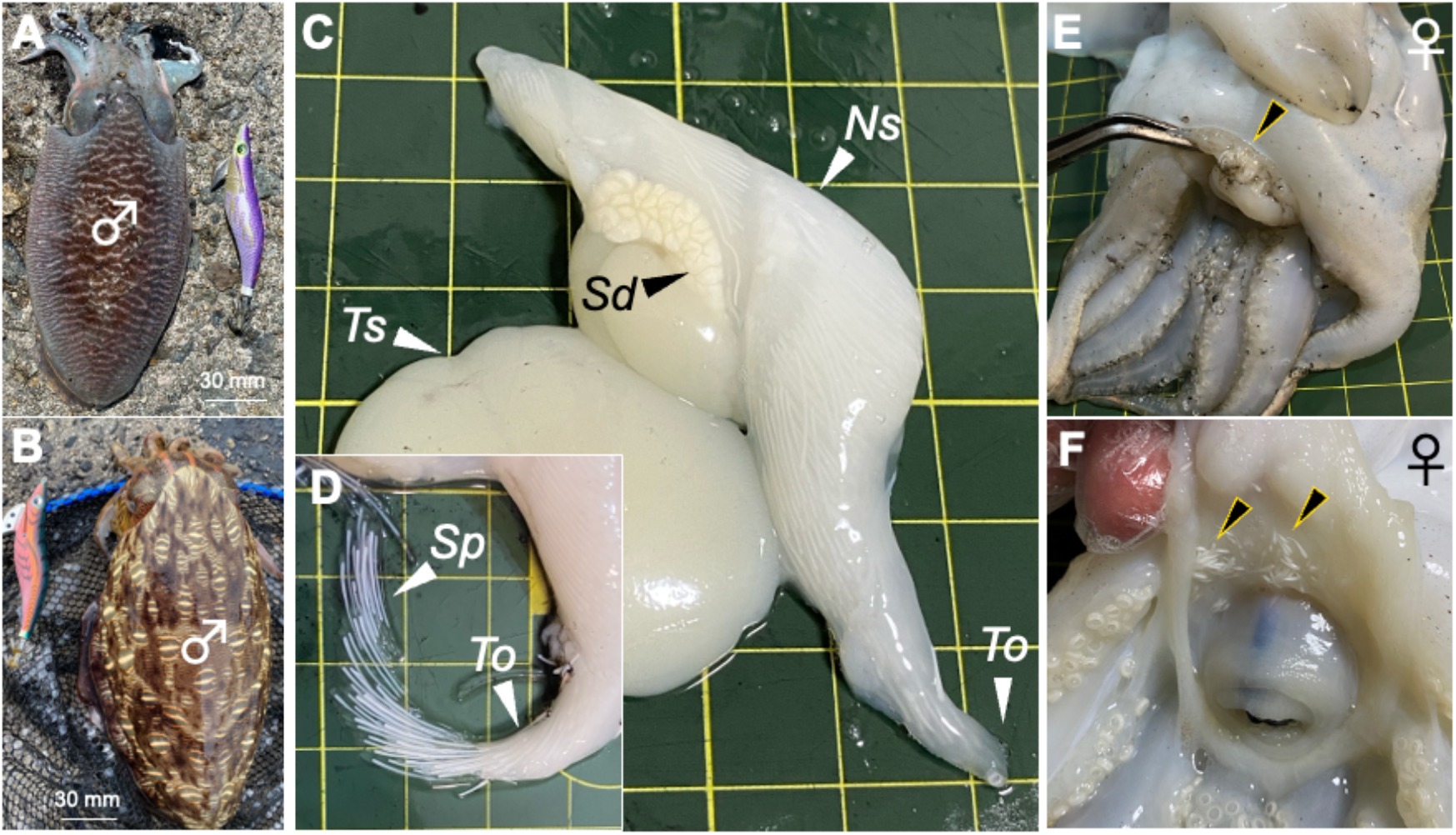
Insemination sites on the female buccal membrane and the male reproductive organ. **A-B**, Males of *S. esculenta* (A) and *S. lycidas* (B) caught by lure fishing. **C**, the isolated male reproductive organ in *S. lycidas. Ts*: testis, *Sd*: spermatophoric duct, *Ns*: Needham’s sac, *To*: terminal organ. **D**, spermatophores expelled by manual compression of the open end of the terminal organ. *Sp*: spermatophores. **E-F**, ventral views of the head region of the female *S. esculenta* (E) and *S. lycidas* (F). *Arrowheads* show spermatangia attached to the ventral side of the buccal membrane. A grid with 10 mm squares.

Measurement of sperm flagellum length (FL) was carried out as described previously (Iwata et al., 2011). Briefly, sperm were released from the spermatophores stored in the terminal organ (Fig. 2E, 2F, To) or from the spermatophoric duct near the testis (Fig. 2E, *Sd*). Sperm were fixed with 4% formaldehyde (Nacalai Tesque, Inc., Kyoto, Japan) in seawater and stored at room temperature. Aliquots of formaldehyde-fixed sperm were mounted on a slide glass and viewed under a microscope (Nikon TE-2000) with a x20 objective lens. Images were captured using a USB camera, and FL was measured using ImageJ software (National Institutes of Health). All analyses were done in RStudio with R 4.4.1 (R Core Team, 2024). A linear model, using the built-in lm function in R, was employed to analyze the relationship between TSI* or GSI* and mantle length as an explanatory variable, yielding the best-fitted regression line and 95% confidence intervals. A generalized linear model from the Gaussian family (function “glm”) was applied to delta sperm length using mantle length, GSI*, or TSI* as explanatory variables.

## Results

All individuals were fully mature (stage IV, Fig. 2A, 2B). Males had spermatophores in Needham’s sac (Fig. 1C, Ns) and sometimes also in the terminal organ (Fig. 2D, To). Females always had spermatangia on the most ventral side of the buccal membrane (Fig. 2E, 2F, arrowheads). GSI* and TSI*, the proxies for maturation and promiscuity, were measured as functions of dorsal mantle length within the population. Both indices showed a decreasing trend with increasing mantle length (LM: *S. esculenta*, GSI*, estimate ± SEM = -0.301 ± 0.049, p < 0.001; TSI*, estimate ± SEM = -0.149 ± 0.027, p < 0.001; *S. lycidas*, TSI*, estimate ± SEM = -0.050 ± 0.014, p < 0.01; Fig. 3). We retrieved sperm from spermatophores situated in the terminal organ and from the spermatophoric duct near the testis, and thereafter compared flagellum length (Fig. 4). In both species, sperm in the terminal organ have longer flagella than those positioned proximal to the testis (Figs. 4A, 4E). In *S. esculenta*, the magnitude of sperm size difference showed a significant decrease as mantle length increases (GLM: estimate ± SEM = -0.118 ± 0.051, p < 0.05, Fig. 4B). In contrast, this difference increased with higher TSI* (GLM: estimate ± SEM = 0.715 ± 0.269, p < 0.01, Fig. 4C) and GSI* values (GLM: estimate ± SEM = 0.356 ± 0.128, p < 0.01, Fig. 4D). In *S. lycidas*, only GSI* exhibited a positive correlation with the magnitude difference in sperm size (estimate ± SEM = 0.540 ± 0.140, p < 0.001, Fig. 4H). These results indicate that in the cuttlefish testis, the developmental program consistently converts sperm from longer to shorter flagella during growth.

**Fig. 3.**
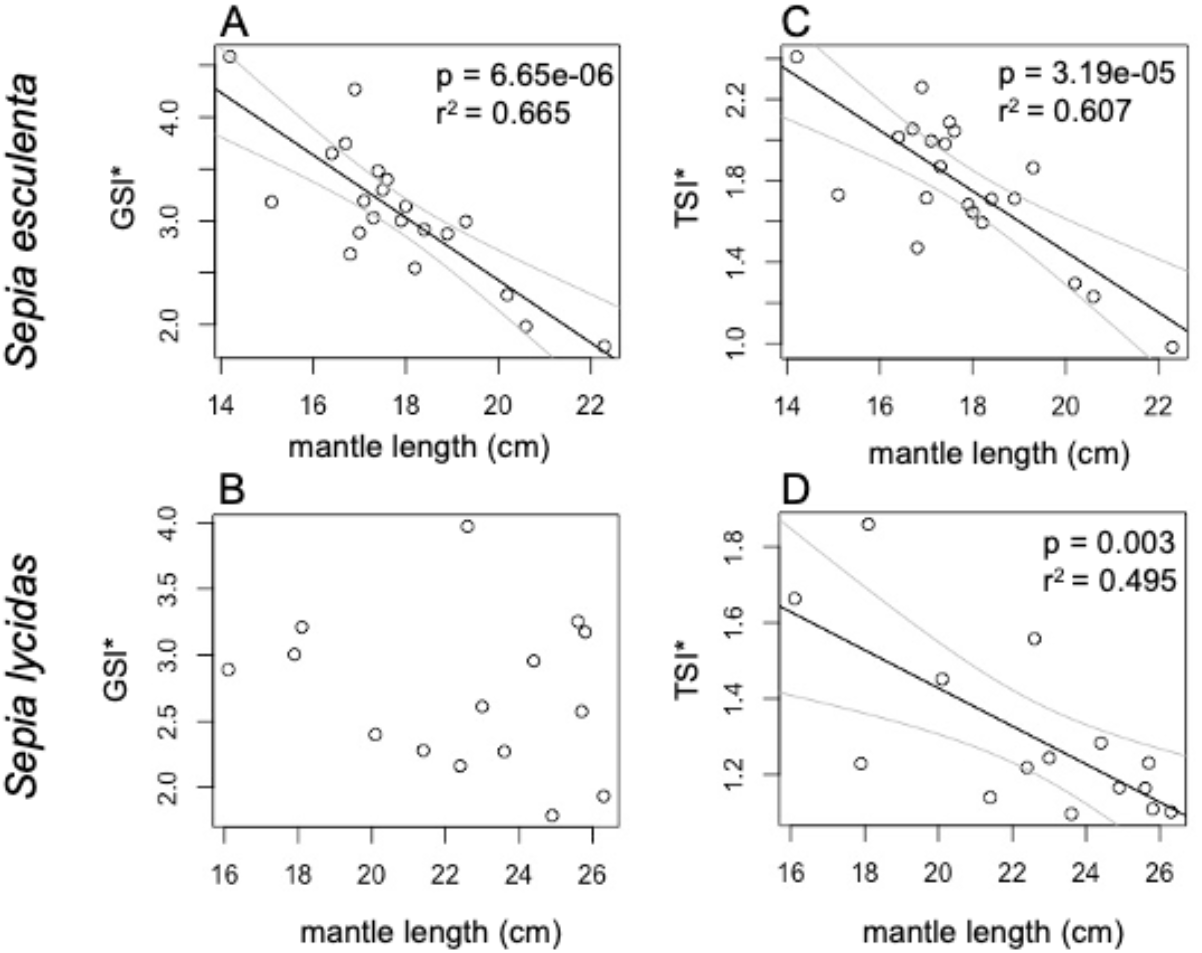
Correlations between somatic growth and gonad growth in two Sepia species. **A-D**, Plots show the relationships between mantle length and gonadosomatic index (GSI*; A, B) or testicular somatic index (TSI*; C, D) in adult males of *S. esculenta* (A, C) and *S. lycidas* (B, D). The indicated lines are a simple linear regression (*solid*) with 95% confidence intervals (*dashed*).

**Fig. 4.**
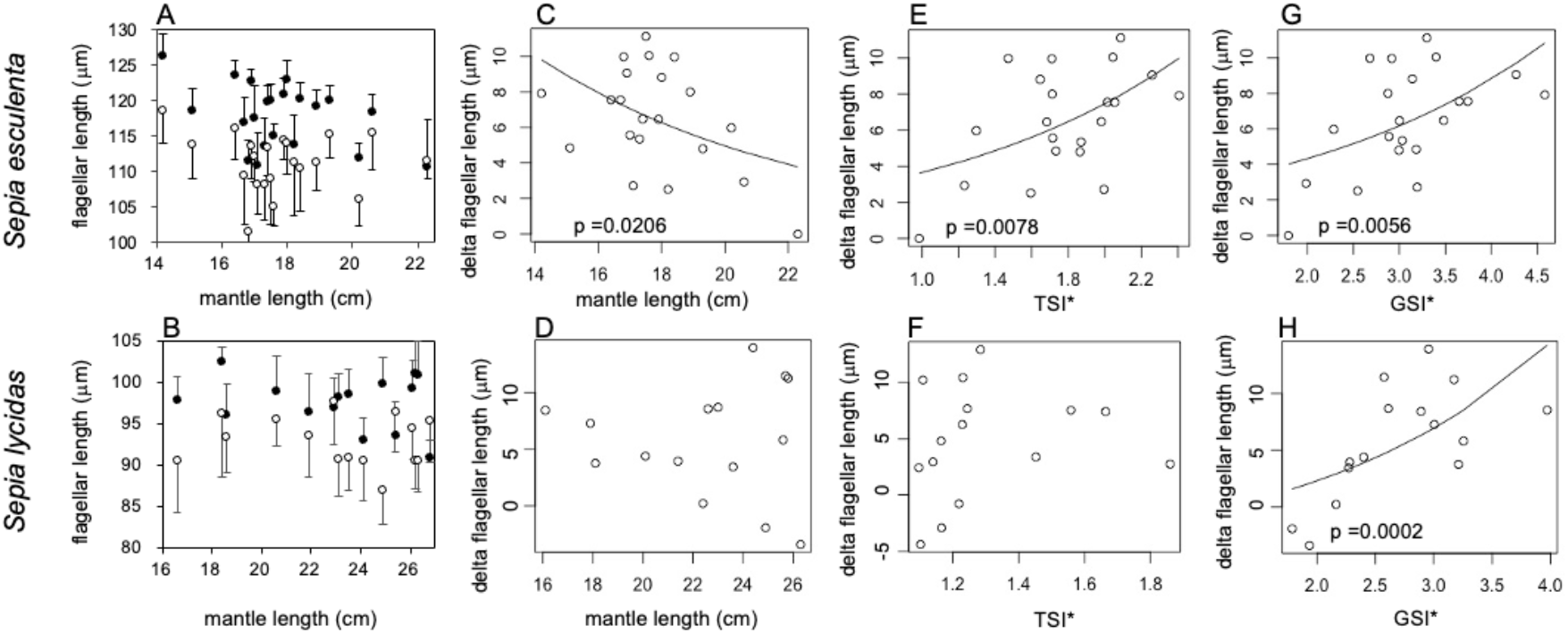
Changes in sperm flagellar length in relation to somatic growth and gonad growth. Panels were arranged to present *S. esculenta* (*the first row*) and *S. lycidas* (*the second row*). **A-B**, plots show the relationships between mantle length and flagellar length of sperm collected from the spermatophore in the terminal organ (*closed circles*) and from the spermatophoric duct near the testis (*open circles*) in *S. esculenta* (A) and *S. lycidas* (B). The mean ± standard deviation of 25-35 collected spermatozoa was plotted. **C-H**, within-male differences in sperm size (delta flagellar length) are plotted against mantle length (C, D), TSI* (E, F), and GSI* (G, H).

## Discussion

The precise length of flagella and cilia in multicellular organisms as well as microorganisms plays fundamental roles in kinematics (Bauer et al., 2021; Ishikawa and Marshall, 2017), physiology (Avasthi and Marshall, 2012; Halte et al., 2025), ecology (Waller et al., 2025) and human health (Chemes et al., 1990; Masyuk et al., 2003; Pazour and Rosenbaum, 2002), and therefore the size is tightly regulated during the processes of generation and regeneration (Marshall, 2023). It is also essential for males to produce sperm with the optimal length of flagellum to achieve both individual and inclusive fertilization success at high rates (Brennen and Winet, 1977; Guasto et al., 2020; Iwata et al., 2011; Kage et al., 2024; Matsuzaki et al., 2021; Reinhardt et al., 2015). For decades, research has been delving into the theoretical modeling and empirical practice of sperm flagellar length being subject to strong postcopulatory sexual selection, i.e., sperm competition and cryptic female choice. Although interspecific variables are significant, general trends indicate that selection favors greater numbers over larger sizes, and that larger/longer sperm are more prevalent in internal fertilizers than in external ones (Kahrl et al., 2021; Wang et al., 2020). This implies that the optimal length of a sperm flagellum is determined by the counterbalance between competitive pressures and environmental factors. These influences operate according to the principles of ejaculate economics, which describes male investment of limited ejaculate resources into each reproductive opportunity (Dowling and Simmons, 2012; Parker and Pizzari, 2010; Xu and Wang, 2014), and sperm ecology, which posits that sperm function is shaped by diverse environments (Guasto et al., 2020; Reinhardt et al., 2015)

We have been studying squid ARTs, in which males produce dimorphic sperm in flagellar length and swimming behaviors (Hirohashi et al., 2013; Iwata et al., 2011). We have long held the hypothesis that sperm dimorphism is linked to different mating positions by males, resulting in two distinct sperm transfer sites on the female. Insemination dimorphism can generate significant differences in post-mating sperm conditions, including the mode of sperm storage, the intensity of sperm competition, and the site of fertilization. We therefore expected that cuttlefish ARTs would not involve bifurcating evolution in the sperm flagellum, since insemination dimorphism is absent. Contrary to our prediction, we found in cuttlefish an intra-male developmental transition in flagellar length from longer to shorter, a phenomenon that we and others have recently observed in the squid *S. lessoniana* (Lin et al., 2019; Tomano et al., 2026). Male *S. lessoniana* undergo a behavioral transition from the sneaker tactic to the consort tactic as they grow, leading to a switch in insemination site (Wada et al., 2005a). Hence, the concurrent changes in sperm size and mating positions further supported our hypothesis (Apostólico and Marian, 2018b; Hirohashi et al., 2021). However, in cuttlefish, it is evident that sperm size can change without switching in mating positions, prompting us to reconsider the set point from which sperm dimorphism has evolved. 1) One potential difference in the fate of sperm is that sneaker sperm, but not consort sperm, are designed to be stored in the female seminal receptacle (Hirohashi et al., 2021). This assumption is based on the squid ARTs, where consorts mate with females immediately before spawning, while sneakers are capable of mating long before spawning (Hirohashi et al., 2016). 2) Another possible explanation is that sneakers invest less ejaculate per mate (Iwata et al., 2011), which may lead to the development of additional traits in sneaker sperm to enhance competition with consort sperm. Supporting this, studies in mammals (Gomendio and Roldan, 1991), birds (Matsuzaki et al., 2021), and fish (Fitzpatrick et al., 2009) have shown that sperm with longer flagella swim faster than those with shorter flagella (but see Kleven et al., 2009). 3) Finally, the relative testis mass (TSI*) has been widely recognized and used as an indicator of promiscuity (Birkhead, 2007; Parker, 2016) (males’ investment in sperm production, but see (Baker et al., 2020)). In cuttlefish, especially in *S. esculenta*, sperm size decreases as TSI* decreases, assuming that larger males should allocate some of their reproductive resources to guarding females, such as increasing body mass. On the other hand, smaller sneakers can invest more reproductive effort in their testes, possibly resulting in the production of larger sperm (Iwata et al., 2021; Ramm and Stockley, 2010). It is also possible that, despite equal opportunity for each spermatozoon to fertilize, the overall fertilization success of sneakers would be lower because consorts predominantly acquire mates. Future studies in cuttlefish ARTs should address the fate of sperm on the female buccal mass as well as the rate of fertilization success for each type of sperm.

Another interesting question is why insemination dimorphism has evolved only in squid, not in cuttlefish. Conversely, male cuttlefish exhibit a unique behavioral trait of sperm removal, a phenomenon not yet observed in squid (Wada et al., 2005b; Wada et al., 2010). Insemination dimorphism and sperm removal may be mutual counterstrategies for second males to mate, thereby increasing their opportunities to sire offspring. Moreover, cuttlefish are masters of sophisticated behaviors, enabling them to mimic females (Hanlon et al., 2005; Norman et al., 1999) and express various courtship displays (Hanlon and McManus, 2020; Nakayama et al., 2024; Schnell et al., 2019), suggesting that pre-ejaculatory sexual selection has operated intensively through these processes to take the lead in mate acquisition. Further comparative approaches with ARTs in squids and cuttlefish will yield better evolutionary insights into cephalopod reproductive strategies.

## Author contributions

KK: data curation, formal analysis, investigation, methodology and writing—review and editing; NH: conceptualization, funding acquisition, data curation, formal analysis, methodology and writing—original draft. All authors gave final approval for publication and agreed to be held accountable for the work performed herein.

## Acknowledgments and funding

We acknowledge the funding support from the Faculty of Life and Environmental Science at Shimane University and Kakenhi (25K02088, 25K09247 and 26K23260 to N.H). This research was conducted in part as part of the SDGs Research Project at Shimane University.

## Declarations

### Ethical statement

The specimens used in this research were obtained under organization approval (Shimane Univ./Dept. Exp. Animals permit #MA2-2, #MA7-09).

